# High-resolution transcriptomic profiling of the aortic cellular landscape during hypertension reveals novel drivers of vascular fibrosis

**DOI:** 10.1101/2025.03.13.642936

**Authors:** Maria Jelinic, Malathi S I Dona, Tayla Gibson Hughes, Gabriella E. Farrugia, Rebecca Harper, Ian Hsu, Asha Haslem, Vivian Tran, Hericka B. Figueiredo Galvao, Henry Diep, Quynh N Dinh, Taylah Gaynor, Tomasz J. Guzik, Matteo Lemoli, Christopher G Sobey, Alex Bobik, Mathew G Lewsey, Alexander R. Pinto, Antony Vinh, Grant R Drummond

## Abstract

**Background:** Aortic stiffening is a consequence of hypertension and a major contributor to end organ damage. A key driver of aortic stiffening is fibrosis involving the excess production of extracellular matrix (ECM) proteins such as collagen, fibronectin and laminin. The present study aimed to identify the cell types and signalling mechanisms that contribute to aortic fibrosis in hypertension.

**Methods and Results:** Male C57BL/6 mice (10-12-week-old) were randomly assigned to a 28-day angiotensin II (0.7 mg/kg/day) or vehicle (saline) infusion via osmotic minipump (*s.c.*). At endpoint, scRNA-seq analysis of 26,196 cells recovered all major aortic cell populations. Among these, fibroblasts exhibited the greatest heterogeneity and shift in gene expression after angiotensin II compared to all other cell types. Gene ontology analyses revealed that after angiotensin II treatment, a particular subcluster of fibroblasts (Fibro-*Cthrc1*) – characterised by its high expression of *Cthrc1* – was especially fibrogenic. Fibro-*Cthrc1* cells were nearly undetectable in aortas from vehicle-infused mice. Transcripts relating to ECM remodelling (*Thbs2*, *Cdh11* and *Postn*) and collagen production (specifically collagen type I, III and V) were more highly enriched in Fibro-*Cthrc1* compared to other fibroblasts within hypertensive aortas. Moreover, GO terms corresponding to profibrotic signalling pathways (i.e., *cell adhesion, extracellular matrix organisation* and *collagen fibril organisation*) were significantly enriched in Fibro-*Cthrc1*. Spatial transcriptomics and immunohistochemistry confirmed the presence of Fibro-*Cthrc1* in the adventitial layer of angiotensin II-infused but not vehicle-infused mice. Finally, analysis of plasma analytes in approximately 24,000 participants of the UK Biobank collection revealed CTHRC1 to be strongly associated with raised systolic blood pressure and pulse pressure, and a strong predictor of the risk of developing hypertension over a 15-year follow-up.

**Conclusion:** Our study identifies a novel fibroblast subcluster, Fibro-*Cthrc1*, as a potential driver of aortic fibrosis and stiffening in hypertension. This cluster is absent in normotensive aortas, suggesting that targeting Fibro-*Cthrc1* therapeutically could prevent aortic fibrosis and its associated hypertensive end-organ damage. Notably, such an approach may avoid compromising physiological extracellular matrix production and vessel integrity.

**Translational perspective:** Aortic stiffening is a hallmark of hypertension resulting from functional (vasoconstriction) and structural (extracellular matrix remodelling) alterations of the vessel wall. While several antihypertensive medications address functional changes, no therapies directly target the causes of the structural remodelling. The therapeutic challenge is to distinguish between physiological and pathological extracellular matrix remodelling. This study identifies a novel highly profibrotic fibroblast cell population (Fibro-*Cthrc1*) present in aortas from hypertensive, but not normotensive mice. This raises the possibility that Fibro-*Cthrc1* may be a key driver of aortic stiffening and a promising future therapeutic target.

## 1. Introduction

Aortic stiffening is a hallmark of hypertension and aging and an independent predictor of all-cause and cardiovascular mortality^1^. Aortic stiffening manifests from changes to the structural (vascular fibrosis and hypertrophy) properties of the vessel wall which impair elasticity^1^. Aortic elasticity is largely governed by the relative amounts of elastin and collagen within the extracellular matrix (ECM) of the vessel wall^2^. A healthy aorta is characterised by its highly developed tunica media, in which elastin fibres are the dominant ECM component and confer elastic properties to the aorta. However, in hypertension, elastin is gradually degraded due to increased pulsatile stress^2^. This is accompanied by a process known as fibrosis, the excess deposition of collagen in the adventitial layer. Collagen is a rigid protein that provides structural integrity to the vessel – allowing it to withstand the increased wall pressures imposed by hypertension but at the expense of elasticity^3^.

While ECM and medial remodelling are characteristic of aortic stiffening, the dynamics of the aortic cellulome—the integrated network of cells that form the aorta—remains unclear. Fibroblasts are traditionally regarded as the key regulators of ECM composition through their high capacity for collagen generation^4^. With regard to several other fibrotic diseases, including cardiac and renal fibrosis, the prevailing paradigm suggests that fibroblasts differentiate into myofibroblasts which have the capacity to produce copious amounts of collagen and thereby drive fibrosis^5^. However, this has never been definitively proven for aortic fibrosis. Single-cell RNA sequencing (scRNA-seq) has provided valuable insights into the cellular heterogeneity of fibroblasts in the aorta in the context of several aortic pathologies, including atherosclerosis and abdominal aortic aneurysms (AAA)^6–8^. While myofibroblasts have been identified by scRNA-seq in the aorta in certain disease states, they are but one of numerous fibroblast subclusters present^7, 9^. Moreover, no studies to date have comprehensively characterised the cellular landscape of the aorta in the context of hypertension and aortic stiffening. Therefore, it remains unclear whether the myofibroblast paradigm is relevant to aortic stiffening in hypertension.

Therefore, to address this gap in knowledge, we performed scRNA-seq and spatial transcriptomics analyses to characterise changes in the aortic cellular landscape during aortic stiffening in angiotensin II-dependent hypertension in mice, with a focus on the aortic fibroblast populations. Our study revealed cell type-specific changes in gene expression during hypertension and aortic stiffening, predominantly in fibroblasts. We confirmed that angiotensin II-induced hypertension is associated with an increase in the number of aortic myofibroblasts. However, even more striking was the appearance of a distinct subpopulation of local fibroblasts that emerged during aortic stiffening in hypertensive mice. This novel fibrogenic cell population is characterised by its exclusive expression of the *collagen triple helix repeat containing-1* (*Cthrc1*) gene. We also characterised key changes in intercellular communication networks and fibroblast differentiation in the context of aortic stiffening. This work provides a valuable resource for vascular cell biology research and offers important insights into the orchestrated cellular and molecular mechanisms driving aortic fibrosis and stiffening.

## 2. Methods

### 2.1 Animals

All animal procedures adhered to guidelines from Directive 2010/63/EU of the European Parliament on the protection of animals used for scientific purposes, the current NIH guidelines, and the National Health and Medical Research Council (NHMRC) of Australia Code for the Care and Use of Animals for Scientific Purposes (8th Edition, 2013). Experiments were approved by the La Trobe University Animal Ethics Committee (AEC# 16-93). Male C57BL/6J mice (10–12 weeks old, 25–30g) were obtained from La Trobe Animal Research and Teaching Facility (Bundoora, Australia). Mice were housed in individually ventilated cages on a 12-hour light/dark cycle at 20°C, with standard food pellets (Barastock, VIC, Australia) and water provided *ad libitum*.

### 2.2 Induction of hypertension and aortic stiffening

Hypertension was induced by subcutaneous implantation of an osmotic minipump (Alzet, USA) delivering angiotensin II (0.7 mg/kg/day) for 28 days. Normotensive control mice received a minipump with 0.9% saline. Blood pressure was monitored via tail-cuff plethysmography (MC4000 Multichannel system, Hatteras Instruments, USA) before and 7, 14, 21, and 28 days after implantation^10^. Aortic stiffening was assessed 28 days post-implantation using high-resolution ultrasound (Vevo 2100, FUJIFILM Visualsonics Inc., Canada) and analyzed with VevoLab and VevoVasc software using the lnD-V loop method^11^.

### 2.3 Aortic single-cell isolation, library preparation and sequencing

At the end of the infusion period, mice were euthanized via CO2 asphyxiation, and blood was collected from the left ventricle. Mice were perfused with PBS containing 5% clexane (400 U/mL) to clear blood from organs. Thoracic and abdominal aortas, with perivascular fat intact and cleaned of lymph nodes, were isolated and placed in ice-cold PBS for further processing. While most previous scRNA-seq studies focused on the inner three vessel wall layers (tunica intima, media, and adventitia)^6, 7, 12^, perivascular adipose tissue (PVAT) was included in this analysis due to its role in vascular homeostasis and disease progression in hypertension. ^13^. The PVAT is also a reservoir for immune cells and fibroblasts^13^ – both of which are involved in vessel remodelling – and hence this layer was included in the analysis (Figure 1A).

**Figure 1.**
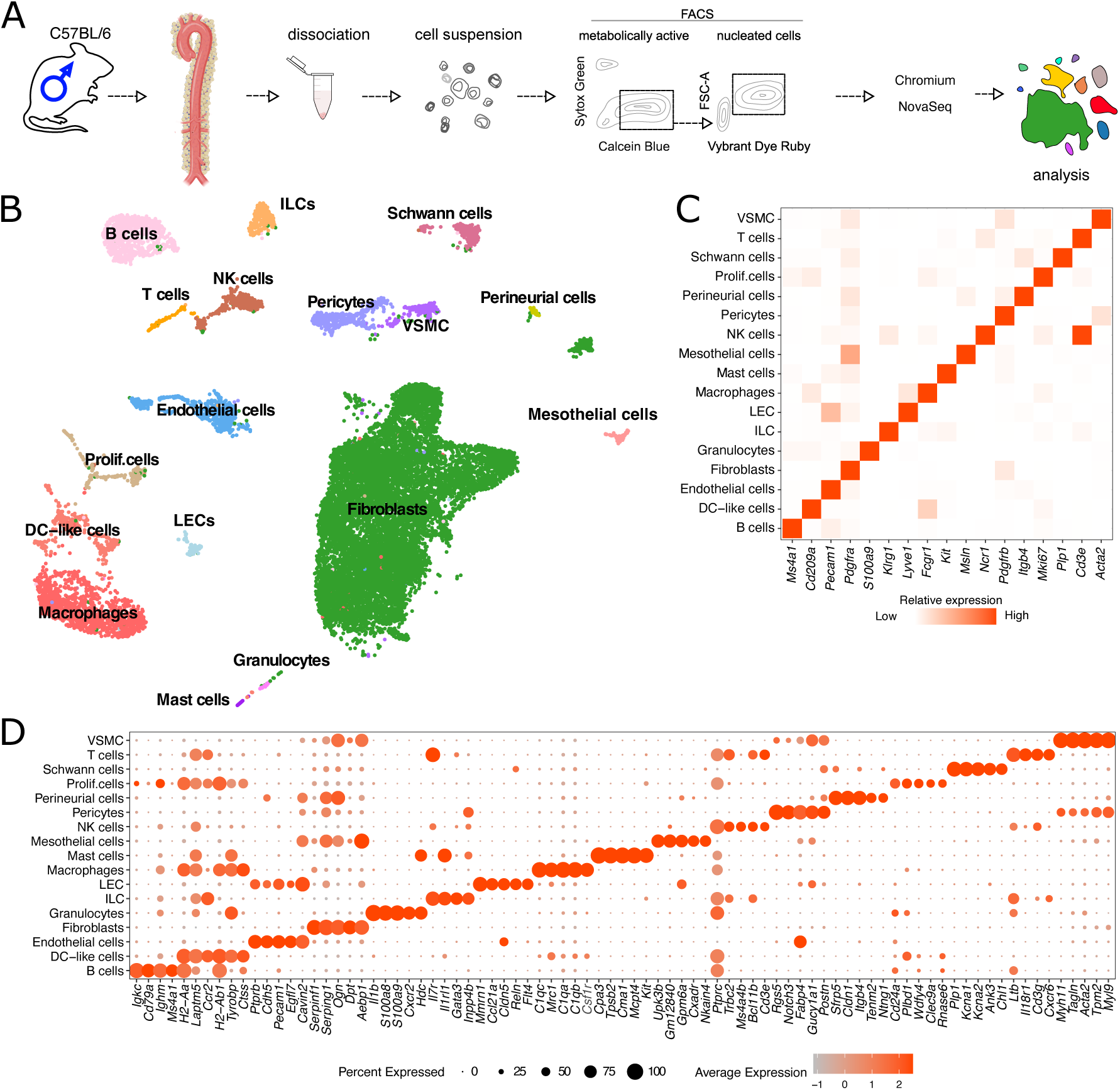
Isolation and analysis of the aortic cellulome by scRNA-seq. Schematic outline of the experimental procedure for the isolation and analysis of mouse aortas by scRNA-seq (A). Uniform Manifold Approximation and Projection (UMAP) of aortic cells analysed by scRNA-seq in vehicle mice. Cells are coloured by distinct cell populations as indicated (B). Heat map of relative expression of canonical cell markers in major aortic cell populations (C). Top 5 distinct genes for each cell population, identified using an unsupervised approach (D). Dot colour and size indicate the relative average expression level and the proportion of cells expressing the gene, respectively, within each cell population (also see Table 1 in the Data Supplement). DC, dendritic cell; FACS, fluorescence-activated cell sorting; FSC-A, forward scatter area; ILC innate-like cell, LEC, lymphatic endothelial cell; NK, natural killer; Prolif, proliferating; and VSMC, vascular smooth muscle cells.

**Table 1.** Characteristics of normotensive patients divided into tertiles of plasma CTHRC1 levels. Data are presented as mean ± SD or proportions (%). BMI, body mass index; SBP, systolic blood pressure; DBP, diastolic blood pressure; HDL, high-density lipoprotein; LDL, low-density lipoprotein; TD index, Townsend deprivation index.

| **CTHRC1 tertile** | **Low**<br>**(n = 4424)** | **Medium**<br>**(n = 4426)** | **High**<br>**(n = 4422)** |
| --- | --- | --- | --- |
| **Age** | 51.56 ± 7.64 | 54.41 ± 7.87 | 56.98 ± 7.73 |
| **Sex (% female)** | 81.7 | 63.2 | 42.4 |
| **Smoking status** |  |  |  |
| **Never (%)** | 59.5 | 58.0 | 54.6 |
| **Former (%)** | 29.0 | 31.0 | 34.2 |
| **Current (%)** | 11.4 | 11.0 | 11.2 |
| **BMI (kg/m<sup>2</sup>)** | 24.94 ± 3.59 | 25.92 ± 3.91 | 27.25 ± 4.36 |
| **SBP (mmHg)** | 121.55 ± 10.30 | 123.98 ± 9.82 | 126.24 ± 9.00 |
| **DBP (mmHg)** | 75.41 ± 7.04 | 76.05 ± 6.95 | 76.81 ± 6.92 |
| **HDL (mmol/L)** | 1.56 ± 0.37 | 1.52 ± 0.38 | 1.44 ± 0.39 |
| **LDL (mmol/L)** | 3.50 ± 0.82 | 3.60 ± 0.82 | 3.63 ± 0.83 |
| **TD index** | -1.58 ± 2.88 | -1.56 ± 2.89 | -1.52 ± 2.97 |
| **Moderate physical activity** |  |  |  |
| **0-2 days/week (%)** | 35.0 | 35.0 | 36.1 |
| **3-4 days/week (%)** | 26.1 | 25.0 | 24.4 |
| **5-7 days/week (%)** | 38.8 | 40.0 | 39.5 |
| **Diabetes prevalence (%)** | 1.2 | 1.5 | 2.6 |
| **Alcohol intake frequency** |  |  |  |
| **Daily (%)** | 17.8 | 19.8 | 20.4 |
| **3-4 days/week (%)** | 26.4 | 24.6 | 23.6 |
| **1-2 days/week (%)** | 29.7 | 27.9 | 27.3 |
| **1-3 days/month (%)** | 11.7 | 11.8 | 11.6 |
| **Occasionally (%)** | 9.8 | 9.4 | 10.3 |
| **Never (%)** | 4.6 | 6.5 | 6.8 |

Aortas were finely minced and transferred to ice-cold enzyme solution (1 mg/mL Dispase II, 4942078001, Roche; 2 mg/mL collagenase IV, LS004188, Worthington Biochem in 0.9 mM CaCl_2_ in PBS), as described previously^14, 15^. Samples were incubated at 37°C for 45 minutes with gentle trituration every 15 minutes. After incubation, samples were placed on ice, filtered through a 70 µm mesh, and centrifuged at 200×g for 15 minutes at 4°C. The supernatant was aspirated, and the pellet re-suspended in 2% FCS, 0.9 mM CaCl2 in PBS. Cells were centrifuged at 400×g for 5 minutes at 4°C, and the supernatant was aspirated. Fluorescence-activated cell sorting (FACS) was performed based on SYTOX Green negative, calcein Blue positive, and Vybrant™ DyeCycle™ positive populations. Sorted cells were counted using a haemocytometer. Approximately 18,000 cells were loaded per channel of Chromium Next GEM Chip G using the Chromium Single Cell 3’ reagent kit (v3.1), and GEMs were created inside the Chromium controller. Library preparation and sequencing were performed using a NovaSeq.

### 2.4 Analysis of single-cell RNA sequencing data

Cell Ranger v3.1.0 (10x Genomics) processed raw sequencing data, aligning reads from FASTQ files to the mm10 transcriptome using the STAR aligner and quantifying transcript expression in each cell. A total of 34,749 aortic cells from normotensive (23,104 cells) and hypertensive (11,645 cells) mice passed initial quality control. scRNA sequencing generated a median of 28,212 reads/cell for vehicle-infused aorta and 33,577 reads/cell for angiotensin II-infused aorta.

Cells expressing < 200 genes or genes expressed in < 3 cells were removed. Cells with > 20% mitochondrial gene expression were filtered out to exclude low-quality and dying cells. The DoubletFinder algorithm was applied to remove potential doublets, selecting only "singlets" for downstream analysis (19,442 vehicle-infused and 10,281 angiotensin II-infused cells). An 8% mitochondrial gene cut-off was applied, leaving 17,571 normotensive and 8,623 hypertensive aortic cells. Normotensive cells were subsampled to 8,623 for balanced analysis. Processed scRNA-seq data were analysed using R v3.6.2, R v4.1.3, and Seurat R package v3.2^16^ and v4.1^17^.

To explore transcriptional heterogeneity and cluster cell types, Principal Component Analysis (PCA) was used for dimensionality reduction, selecting 40 principal components (PCs). PC loadings were used for graph-based clustering at a resolution of 1.2 and for visualization with Uniform Manifold Approximation and Projection (UMAP). Cell clusters were annotated based on known marker genes. Figures were generated using Seurat and ggplot2 R packages^18^.

### 2.5 Differential expression analysis

To identify differentially expressed genes, we filtered genes expressed in at least 10% of cells in one of the groups. The MAST method was used for differential expression analysis, considering cellular detection rate as a covariate^19, 20^. An uncorrected *P* < 0.01 threshold defined statistical significance.

### 2.6 Gene ontology analysis

Gene ontology (GO) over-representation analysis for differentially expressed gene lists (uncorrected *P* <0.01) was performed using the enrichGO function from clusterProfiler R package version 4.2.2^21^. All mappings were based on the data from *org.Mm.eg.db*: Genome wide annotation for mouse^22^, R package version 3.14.0, which primarily uses Entrez Gene identifiers and gene ontology information from http://geneontology.org for GO mapping. The over-representation of GO Biological Process terms was calculated using the list of genes identified in the experiment as the background gene list for *Mus musculus* with gene set sizes between 10-500 genes. GO Biological Process terms that have semantic similarity higher than the cut-off of 0.7 were treated as redundant terms and discarded using simplify function from the clusterProfiler R package. The Benjamini-Hochberg adjusted P-value cut-off of *P* < 0.05 was used to determine statistically significant GO Biological Process terms.

### 2.7 Ligand-receptor intercellular communication analysis

CellChat R package version 1.5.0^23^, was used to infer the intercellular communication pathways between each aortic cell population to implement the signalling pathway networks visualizations. Only ligands and receptors with non-zero expression in > 10% of cells in a particular cell population were considered an “expressed” ligand/receptor interaction. To construct the potential cell-to-cell communication network, expressed ligands were linked with their corresponding receptors between and within the major aortic cell populations. Gene ontology analyses were performed on significantly up and down regulated ligand/receptor gene lists in angiotensin II-infused mouse aorta to identify their functional enrichments.

### 2.8 Trajectory analysis

RNA trajectory analysis was performed on fibroblast cells using Monocle3^24–26^ as described in GitHub page https://github.com/KramannLab/Murine_heart_map^27^. A specific trajectory, starting from Fibro9 to Fibro-*Cthrc1* was selected for further examination of the differentiation of Fibro-*Cthrc1* cells. The pseudotime, a measure of progression over trajectory, was calculated for the selected Fibro-*Cthrc1* trajectory starting at Fibro9 sub population. To identify genes with comparable expression patterns over pseudotime, the top 500 significantly differentially expressed genes across the selected trajectory were combined and subjected to hierarchical clustering. The resulting gene clusters were then visualized on a heatmap. Furthermore, gene ontology analysis was conducted to determine enriched GO terms associated with each gene cluster. The smooth functions were employed to plot the gene expression measures along pseudotime, utilizing the "geom_smooth" function from the "ggplot2" package. The method used for smoothing was the generalized additive model, with the default formula and a negative binomial distribution.

### 2.9 Histopathology and immunohistochemistry

Freshly isolated thoracic and abdominal aortas from vehicle- and angiotensin II-infused mice were formalin-fixed, paraffin-embedded, and sectioned (5 μm). Aortic adventitial collagen content was quantified using Masson’s trichrome or picrosirius red staining^11^. with percentage area assessed in a blinded fashion using ImageJ software. To validate CTHRC1 protein presence, immunohistochemistry was performed on aortic tissue. Paraffin-embedded slides were dewaxed, underwent antigen retrieval in citrate buffer (pH 6.0) at 100°C for 11 minutes, cooled, washed, and blocked. CTHRC1 primary antibody (Abcam, ab85739) was diluted 1:1000, with rabbit IgG isoform (Invitrogen, 08-6199) as a negative control, and incubated at 4°C overnight. Secondary antibody detection used MACH 4 Universal HRP-polymer, betazoid DAB-chromagen, and DAB Sparkle as per manufacturer’s instructions. Sections were dehydrated, mounted with DPX mounting media, and cover slipped.

### 2.10 Spatial transcriptomics expression assay and analysis

Following tissue permeabilization optimization, fresh-frozen aortas were embedded in OCT (Tissue-Tek) and cryosectioned into 10 μm sections. Sections were placed on pre-chilled Visium Spatial Gene Expression Slides (10x Genomics), stained with haematoxylin and eosin, and imaged using an Olympus BX50 microscope with Mosaic Software v2.4. The spatial transcriptomics assay was performed as per the Visium user guide. After quality control via TapeStation (Agilent), library preparation was done at the La Trobe Genomics Platform, and sequencing was performed using a NovaSeq6000. Mean reads per spot ranged from 143,000 to 336,000, aligned to the mm10 mouse reference genome.

Raw sequencing files were pre-processed in SpaceRanger (10x Genomics, v2.1), and H&E images were aligned in Loupe Browser (10x Genomics) with barcodes. Analysis was conducted using RStudio, with filtered feature-barcode expression matrices from SpaceRanger inputted into the Seurat suite (v5.0.1)^28^. Individual count matrices were merged and normalized using SCTransform^29^, followed by additional normalization with Harmony for batch effect correction^30^. Spots with <200 UMIs were filtered out. The aortic cellulome dataset was integrated with the merged spatial transcriptomic object and used as a reference for major cell types.

### 2.11 UK Biobank data analysis

Human data used in this study are available via application to the UK Biobank under Application Number 93156. The UK Biobank is a population-based cohort of ∼500,000 participants aged 40-69 years, recruited between 2006-2010^31^. Participant data included electronic health records, biomarkers, genetic data, imaging data, and physical measurements. Participants provided written informed consent. The UK Biobank Pharma Proteomics Project measured 2,941 blood plasma analytes in 54,219 participants using the antibody-based proximity extension assay^32^. Prospective analyses were conducted in 13,272 normotensive participants (not on BP-lowering medications, SBP <140 mmHg, DBP <90 mmHg, no hypertension diagnosis). Participants were divided into tertiles based on baseline plasma CTHRC1 levels (Table 1). Kaplan-Meier curves for incident hypertension were calculated for each tertile, and Cox regression determined the hazard ratio for developing hypertension (ICD10 code I10). General linear model analyses in 23,564 subjects (not on BP-lowering medications) assessed associations between plasma CTHRC1 and blood pressure indices (SBP, DBP, PP), adjusting for age, sex, BMI, smoking status, and alcohol intake frequency.

## 3. Results

### 3.1 Single-cell transcriptomic profiling of the healthy mouse aorta

Our study aimed to understand how cellular activity and function change during aortic stiffening in hypertension using scRNA-seq of aortas from mice treated with vehicle or angiotensin II (Figure 1A). To ensure high quality genomic data, only live, metabolically active, nucleated cells (Sytox Green^-^/Calcein Blue^+^/Vybrant Dye Ruby^+^) were isolated for scRNA-seq analyses (Figure 1A, Supplementary methods)^15^. A total of 17,571 aortic cells from normotensive mice were assessed using scRNA-seq (Figure 1B). Examination of canonical genes corresponding to various cell types and genes enriched in cell clusters revealed a diverse aortic mouse cellulome comprising at least 17 distinct cell classes (Figure 1B-D, Supplementary Figure 1). These included previously well characterised cell types of the aorta such as fibroblasts (*Pdgrfa, Col1a1*), VSMCs (*Acta2, Tagln, Mhy11*), pericytes (*Vtn*, *Pdgrfb*), endothelial cells (*Pecam1*), as well as immune cell populations including B cells (*Ms4a1*), T cells (*Cd3e*), macrophages (*Fcgr1*), dendritic cell (DC)-like cells (*Cd209e*), mast cells (*Kit*), natural killer cells (*Ncr1*) and granulocytes (*S100a9*)). Additionally, cell types not normally associated with the vascular system were also identified including Schwann cells (*Plp1*), perineurial cells (*Itgb4*), mesothelial cells (*Msln*), innate lymphoid cells (ILC; *Klrg1*), and lymphatic endothelial cells (LEC; *Lyve1;* Figure 1C-D, Supplementary Table 1).

### 3.2 Aortic cellular heterogeneity during hypertension and aortic stiffening

To profile the dynamics of cellular abundance and gene expression, we compared scRNA-seq data from aortas from angiotensin II-infused mice, with those from vehicle-infused mice (Figure 2A). Angiotensin II increased systolic blood pressure by approximately 40 mmHg from day 7 of infusion (Figure 2B). At the day 28 endpoint, aortic stiffening and fibrosis were confirmed in angiotensin II-infused mice by higher pulse wave velocity, increased collagen fibre deposition, and expansion of the adventitial layer compared with normotensive mice (Figure 2C and D). By combining transcriptomic data from normotensive and hypertensive mouse aortas, we obtained a total of 26,194 cells and identified an even more diverse array of cell types and subclusters (Figure 2E, Supplementary Table 2). Examination of cell abundances within this scRNA-seq dataset revealed several cell populations that changed in relative abundance in response to angiotensin II, including one cell subcluster that was only present in aortas from angiotensin II-treated mice (Supplementary Figure 1).

**Figure 2.**
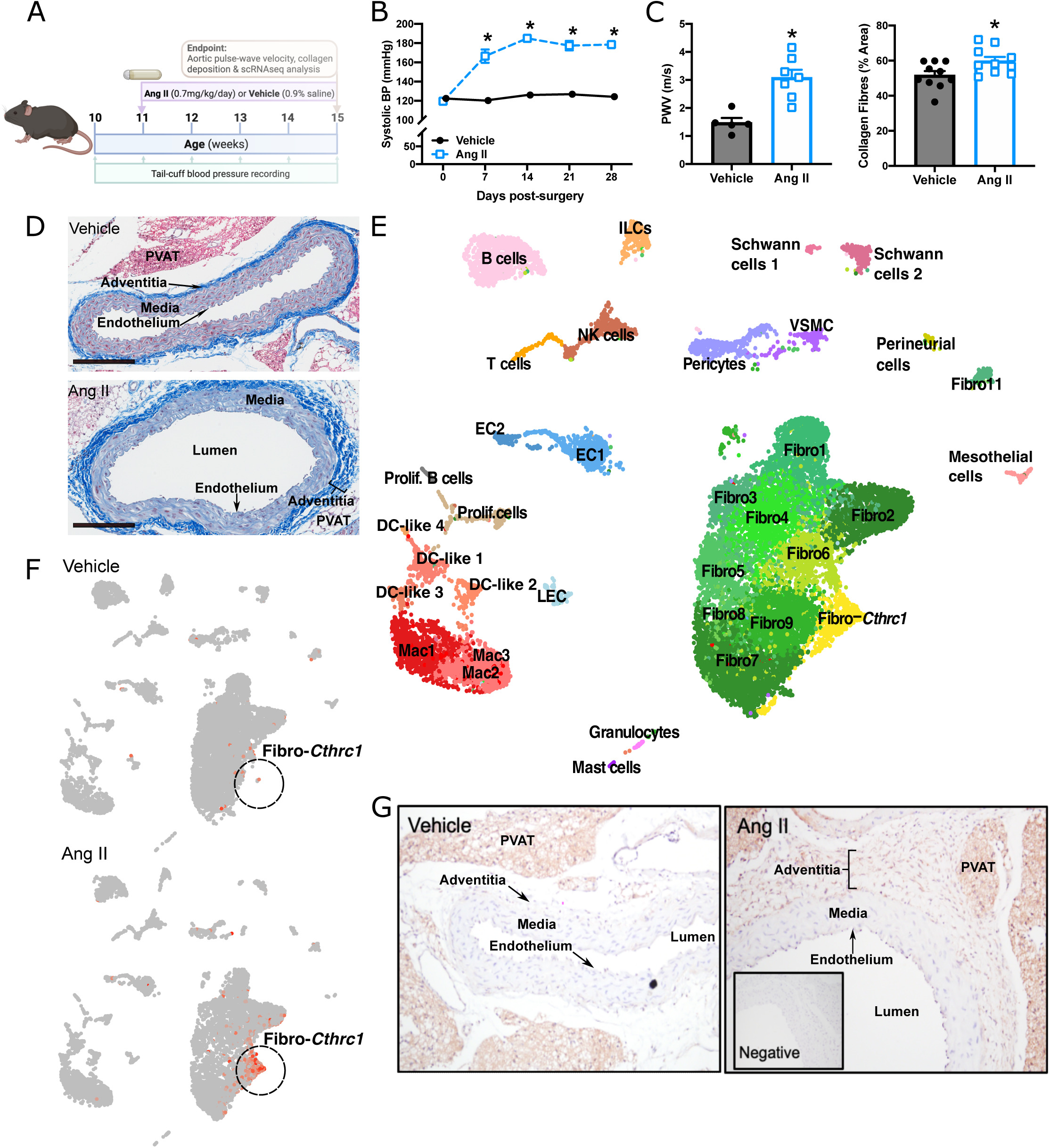
Histological and cellular changes in the aorta after induction of hypertension and aortic stiffening. Experimental schema for angiotensin II (Ang II) induction of hypertension and aortic stiffening. (A) Changes in systolic blood pressure (BP; B), pulse-wave velocity (C-left) and aortic collagen content (C-right) in vehicle (black closed circles) and Ang II-treated male mice (blue open squares; n = 3/group). Representative micrograph of Masson’s trichrome-stained aortic sections from mice infused with vehicle- or Ang II (D). UMAP of aortic cells analysed by scRNA-seq in both vehicle- and Ang II-treated mice (E). Each dot represents a cell with distinct cell clusters and subclusters labelled. Feature UMAP of aortic cells isolated from vehicle- and Ang II-treated (F) mice. Each dot represents a cell, and cells are coloured in red relative to *Cthrc1* gene expression (red = high; grey = low/no gene expression). Representative images of immunohistochemical staining for collagen triple-repeat helix containing protein 1 (CTHRC1) in aortas from vehicle- and Ang II-treated mice (G). Brown staining is indictive of CTHRC1 expression. DC, dendritic cell; ILC innate-like cell, Fibro, fibroblast; LEC, lymphatic endothelial cell; NK, natural killer; Prolif, proliferating; and VSMC, vascular smooth muscle cells. \**P* < 0.05, Micrograph scale bars = 200 μm.

The most striking difference between the two groups was the presence of a prominent fibroblast subcluster in aortas from angiotensin II-infused mice, which was virtually absent in aortas from vehicle-infused mice (Figure 2F, Supplementary Table3). The top uniquely expressed genes in this hypertension-specific fibroblast subcluster were *Cthrc1*, *Comp*, *Col11a1*, *Col8a2* and *Thbs2* (Supplementary Figure 2; subcluster named Fibro-*Cthrc1* hereon). These data were further validated via immunohistochemistry studies. CTHRC1^+^ cells were localised in the aortic adventitia of angiotensin II-infused mice but absent in this layer in vehicle-infused mice (Figure 2G). CTHRC1^+^ staining was also detected in the PVAT, but there were no apparent differences between the angiotensin II- and vehicle-infused groups.

### 3.3 Cell-specific changes in gene expression during hypertension and aortic stiffening

Angiotensin II infusion altered gene expression in several aortic cell populations. Fibroblasts were by noticeably the most affected with 1000 upregulated and 1681 downregulated genes. They were followed by macrophages (790 upregulated; 482 downregulated), with endothelial cells, proliferating cells, B cells and dendritic cells all displaying a similar degree of differentially expressed genes (144-352 upregulated; 200-260 downregulated; Figure 3A, Supplementary Table 4). The top upregulated genes following angiotensin II infusion followed largely cell-specific patterns, although immunoglobulin genes *Ighm, Igkc* and *Jchain* were upregulated in multiple cell types (Figure 3B, Supplementary Table 4). Of the other prominently upregulated genes, several have previously been implicated in hypertension and vascular stiffening such as *Vcam1* (upregulated here in mesothelial cells)^33^, *Tspan13* (in endothelial cells and macrophages)^34^, *Rgs19* (in B cells)^35^, and *Reln* (in Schwann cells)^36^ (Figure 3B). Focusing on fibroblasts, genes associated with ECM remodelling (*Cthrc1*, *Comp*, *Col8a1*, and *Col11a1*) were among the most highly upregulated (Figure 3B).

**Figure 3.**
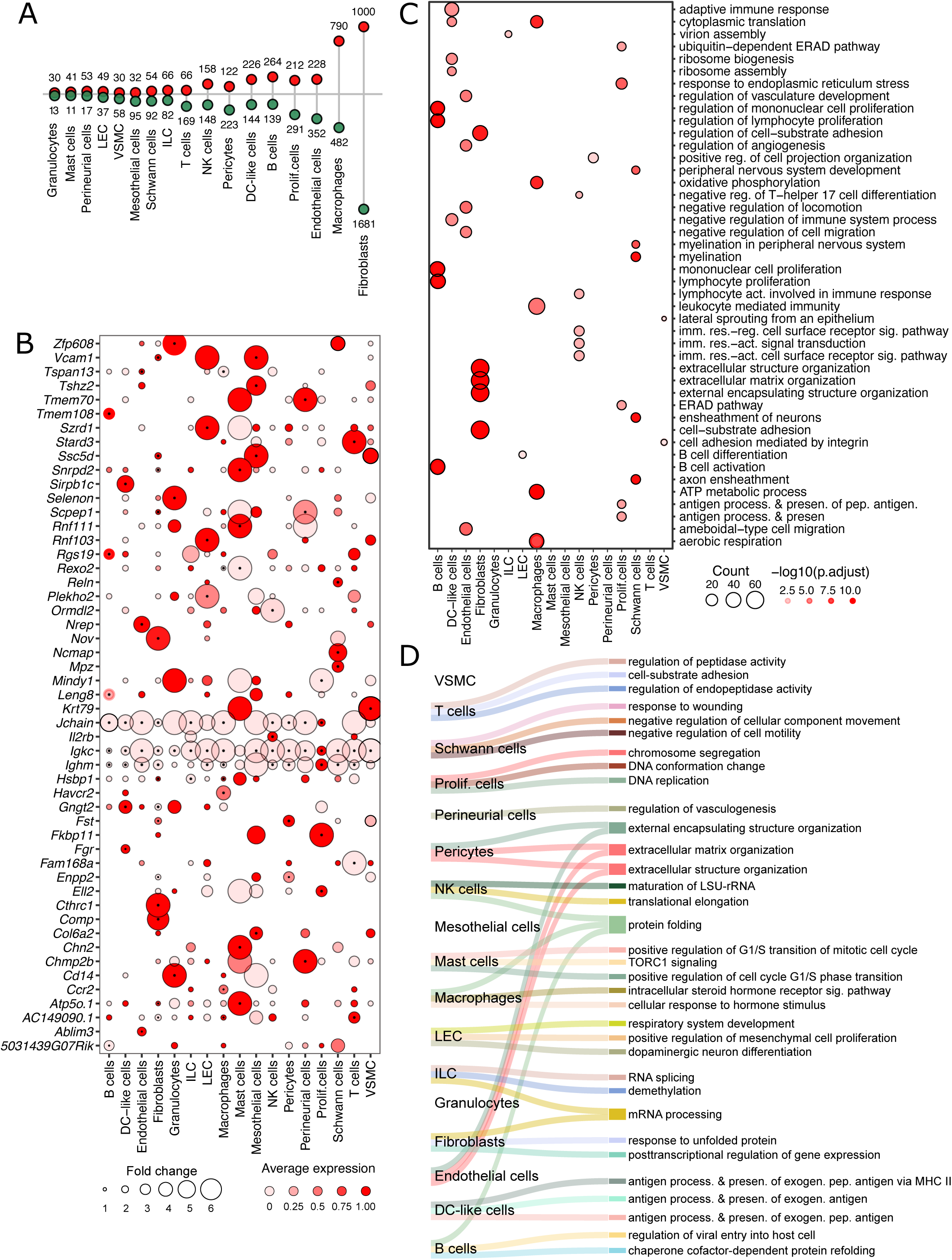
Gene expression changes in aortic cell populations. Lollipop plot summarizing number of up- and down-regulated genes (uncorrected *P* < 0.01) in major aortic cell types from angiotensin II-(Ang II) treated mice, relative to vehicle-treated mice (red = up regulated, green = down regulated; A). Dot plot summarizing the expression of genes identified as within the top 10 genes upregulated in response to Ang II administration, for each cell type (B). Dot colour and size are proportional to average expression within each cell cluster in Ang II cells and fold increase in Ang II cells relative to control cells, respectively. The black dots in the centre of some circles indicate the statistical significance of *P* < 0.01. Dot plot summarizing top 5 gene ontology (GO) terms up regulated in aortic cell populations with corrected *P*<0.05 (C). Dot colour and size proportional to the statistical significance and number of up regulated genes enriched in each GO term, respectively (also see Supplementary Table 5). Sankey plot summarizing the top 3 GO terms for down regulated genes of each cell type with corrected *P*<0.05 (D). Connections indicate GO terms associated with each cell type (also see Supplementary Table 6). Act, activating; DC, dendritic cell; imm, immune; ILC innate-like cell, LEC, lymphatic endothelial cell; NK, natural killer; pep, peptide; presen, presentation; Prolif, proliferating; reg, regulation; VSMC, vascular smooth muscle cells; Neg., negative; reg., regulation; res., response; and sig, signalling

Examination of top five GO terms enriched in upregulated genes revealed concerted gene expression programs occurring in multiple cell types (Figure 3C, Supplementary Table 4). For example, GO pathways associated with immunity and inflammation such as *‘adaptive immune response’*, *‘regulation of lymphocyte proliferation’*, *‘leukocyte cell adhesion’*, *‘antigen processing and presentation’* and *‘leukocyte mediated immunity’* were upregulated in immune cell populations (B cells, DC-like cells and macrophages). Meanwhile, fibroblasts were enriched for terms associated with ECM remodelling and fibrosis such as *‘extracellular matrix organisation’* and *‘extracellular structure organisation’* (Figure 3C). Many biological processes were also downregulated in aortic cell types after angiotensin II infusion (Figure 3D; Supplementary Figure 3B-D). For example, in stark contrast to the changes observed in fibroblasts, collagen genes (*Col1a1*, *Col1a2* and *Col3a1*) were downregulated in multiple cell types (B cells, DC-like cells, endothelial cells, macrophages, NK cells, pericytes, proliferating cells and T cells; Supplementary Figure 3A). Likewise, genes correlating to processes associated with *‘extracellular matrix organization’* and *‘extracellular structure organization’* were downregulated in pericytes and endothelial cells (Figure 3D; Supplementary Figure 3D), while GO terms relating to *‘antigen processing and presentation’* were downregulated in DC-like cells. The most frequently downregulated gene sets among aortic cell types were those aligning to the *‘protein folding’* GO term (i.e. in NK cells, macrophages and B cells; Supplementary Table 6). This is suggestive of an unfolded protein response in these cell types.

### 3.4 Hypertension and aortic stiffening increases aortic paracrine signalling networks

In addition to the cell-specific changes described above, we hypothesised that angiotensin II infusion would also alter aortic paracrine signalling networks between cells. To test for this, we mapped ligands and their cognate receptors in all aortic cell populations, in both normotensive and hypertensive mice. In normotensive mice, fibroblasts, VSMCs and Schwann cells showed the greatest number of incoming and outgoing intercellular ligand-receptor interactions (Figure 4A-B, Supplementary Figure 4). Angiotensin II infusion increased the number of intercellular connections in most cell types, but most strikingly in fibroblasts, VSMCs, LECs, mesothelial cells and perineurial cells (Figure 4B, Supplementary Table 7). Interestingly, compared to vehicle-treated mouse aortas (Figure 4C), the incoming paracrine interactions were increased in angiotensin II infusion in all cell types (particularly VSMCs, macrophages, granulocytes and LECs) (Figure 4D). However, the number of outgoing signals were only increased in fibroblasts, VSMCs, perineurial cells, mesothelial cells and pericytes (Figure 4C-D).

**Figure 4.**
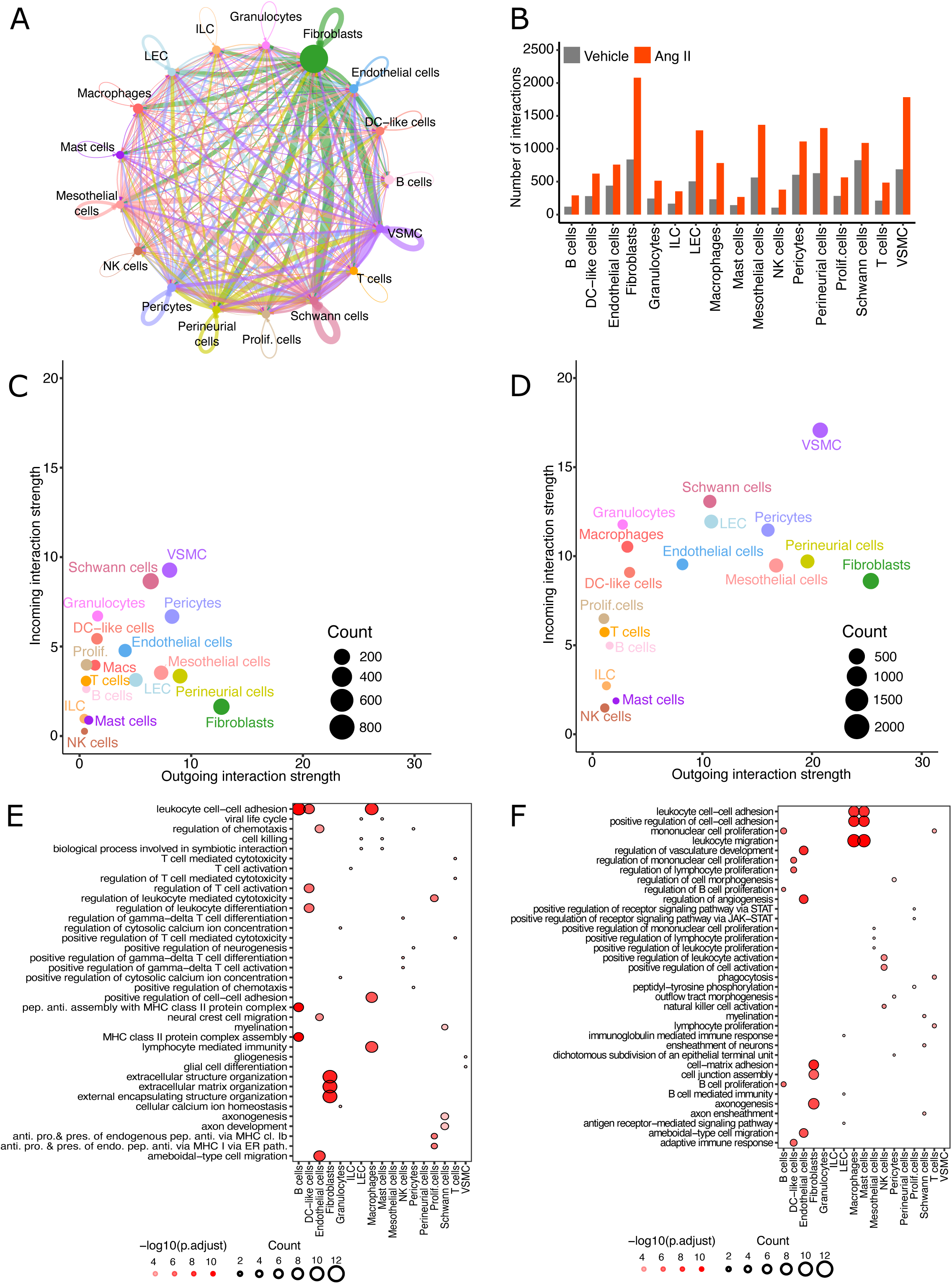
Intercellular communication network of the aortic cellulome. Cell-chat cell-cell communication network summarizing putative intercellular communication network between different aortic cell types from vehicle-treated male mice (A). Lines represent potential interconnections between cell types, with line thickness proportional to the number of ligand-receptor interactions between cell types and line colour reflecting the cell population producing transcript coding for the ligand. Bar plot summarizing the total number of signals transmitted and received from each aortic cell population (grey-vehicle, red-Ang II B) Scatter plot summarizes outgoing and incoming signal strengths for each cell population in vehicle-(C) and Ang II-treated (D) mice. Dot plot summarizing top 3 GO terms enriched in a set of genes that encode ligands (E), and receptors (F) upregulated after Ang II treatment. GO terms are ordered by their frequency of significant enrichment in different aortic cell populations. Dot colour and size proportional to the statistical significance and number of up regulated genes enriched in each GO term, respectively. Anti, antibody; cl, class; DC, dendritic cell; endo; endogenous; ILC innate-like cell, LEC, lymphatic endothelial cell; NK, natural killer; path, pathway; pep, peptide; presen, presentation; pro, production; Prolif, proliferating; VSMC, vascular smooth muscle cells.

To better understand these changes, the top GO terms were determined separately for upregulated genes encoding ligands and receptors for each cell type. Most GO terms corresponding to ligand genes were associated with inflammation in leukocyte cell types (predominantly B cells, DC-like cells and macrophages) and ECM remodelling in fibroblasts (Figure 4E, Supplementary Table 9). Similarly, the top upregulated GO terms for receptor genes were associated with inflammation in leukocyte cell types (predominantly B cells, macrophages and mast cells; Figure 4F, Supplementary Table 10). GO terms associated with downregulated ligand genes included *collagen fibril organization* in B cells, NK cells, proliferating cells and T cells (Supplementary Figure 5A, Supplementary Table 11). Furthermore, downregulated receptor coding genes from B cells were associated with GO processes related to calcium signalling in LECs and pericytes; and inflammation in DC-like cells (Supplementary Figure 5B, Supplementary Table 12).

### 3.5 Elevated fibroblast signalling activity in the hypertensive mouse aorta

Our intercellular communication analyses showed that angiotensin II infusion augmented several interactions in most aortic cell types. To better characterise these interactions, we further mapped the intercellular circuits that were strengthened or induced by angiotensin II. Notably, the number of incoming and outgoing interactions increased most in fibroblasts and VSMCs (Figure 5A; bar plots). While angiotensin II treatment increased the number of incoming/outgoing signals of fibroblasts and VSMCs from/to all major cell types, the most notable changes were in fibroblast-fibroblast and fibroblast-VSMC communications (Figure 5A; heatmap).

**Figure 5.**
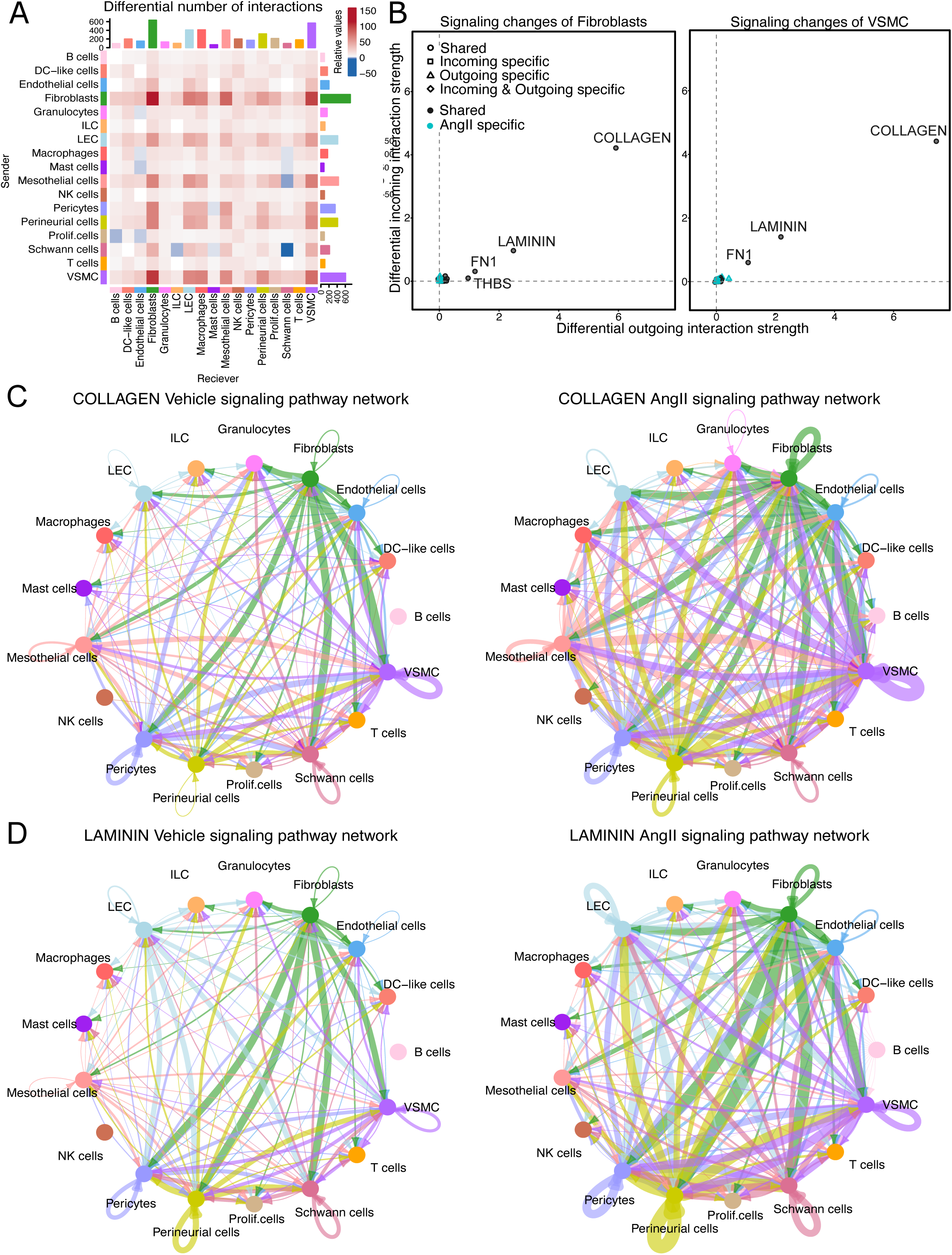
Changes signalling pathway networks of the aortic cellulome. Heatmap showing the differential number of interactions between cell population in the Ang II aortic signalling pathway network compared to vehicle (A). Coloured squares corresponding to the relative changes of the number of interactions (blue = decreased, red = increased). The top-coloured bar plot represents the sum of column values displayed in the heatmap (incoming signals) and the right coloured bar plot represents the sum of the row values (outgoing signals) for each cell type. Bar plot colours reflect the different aortic cell populations. Scatter plots summarizing the specific signalling changes of fibroblasts and VSMC between vehicle and Ang II signalling networks (B). Network graphs visualizing inferred communication network of COLLAGEN signalling pathway in vehicle- and Ang II-treated mice (C). Edge colours are consistent with the sources as sender, and edge weights are proportional to the interaction strength. Thicker edge line indicates a stronger signal. Network graphs visualizing inferred communication network of LAMININ signalling pathway vehicle- and Ang II-treated mice (D). DC, dendritic cell; ILC innate-like cells, LEC, lymphatic endothelial cells; NK, natural killer; prolif, proliferating; and VSMC, vascular smooth muscle cells.

To better understand these changes, the strength of outgoing and incoming interaction signalling pathways were assessed within fibroblasts and VSMCs. Signalling for the ECM proteins, collagen and laminin, were increased to the greatest extent by angiotensin II infusion in fibroblasts and VSMCs (Figure 5B-D). Fibronectin 1 (*Fn1*) and thrombospondin signalling (also ECM proteins) were also increased in fibroblasts from hypertensive mice. Interestingly, laminin signalling was also profoundly increased by angiotensin II in perineurial cells, pericytes, LECs and Schwann cells. Overall, these data suggest fibroblasts and VSMCs are the most profibrotic cell types in the aorta following angiotensin II infusion.

### 3.6 Fibro-Cthrc1 are highly profibrotic

Our scRNA-seq analysis defined 11 fibroblast subclusters (Figure 6A). Given ECM production is primarily mediated by fibroblasts, we sought to determine which of these subclusters are likely to be the predominant contributors to fibrosis. Several fibroblast subclusters changed in relative abundance in response to angiotensin II. Fibro2, Fibro7 and, most notably, Fibro-*Cthrc1* subclusters were expanded (∼1-, ∼2- and ∼96-fold increase, respectively) in the aorta after angiotensin II infusion, whereas Fibro3, Fibro4, Fibro5 and Fibro9 were reduced (Figure 6B, Supplementary Table 3). Given the magnitude of Fibro-*Cthrc1* expansion after angiotensin II infusion, we next examined for the possible origin of this subcluster. Cells corresponding to Fibro-*Cthrc1* subcluster were very low in number in vehicle-infused mice. Given that no proliferative fibroblast subpopulations were observed (assessed by expression of proliferative markers *Top2* and *Mki67*; Supplementary Figure 6), it is conceivable that fibroblast differentiation is the major process contributing to the expansion of the Fibro-*Cthrc1* subcluster during the development of hypertension.

**Figure 6.**
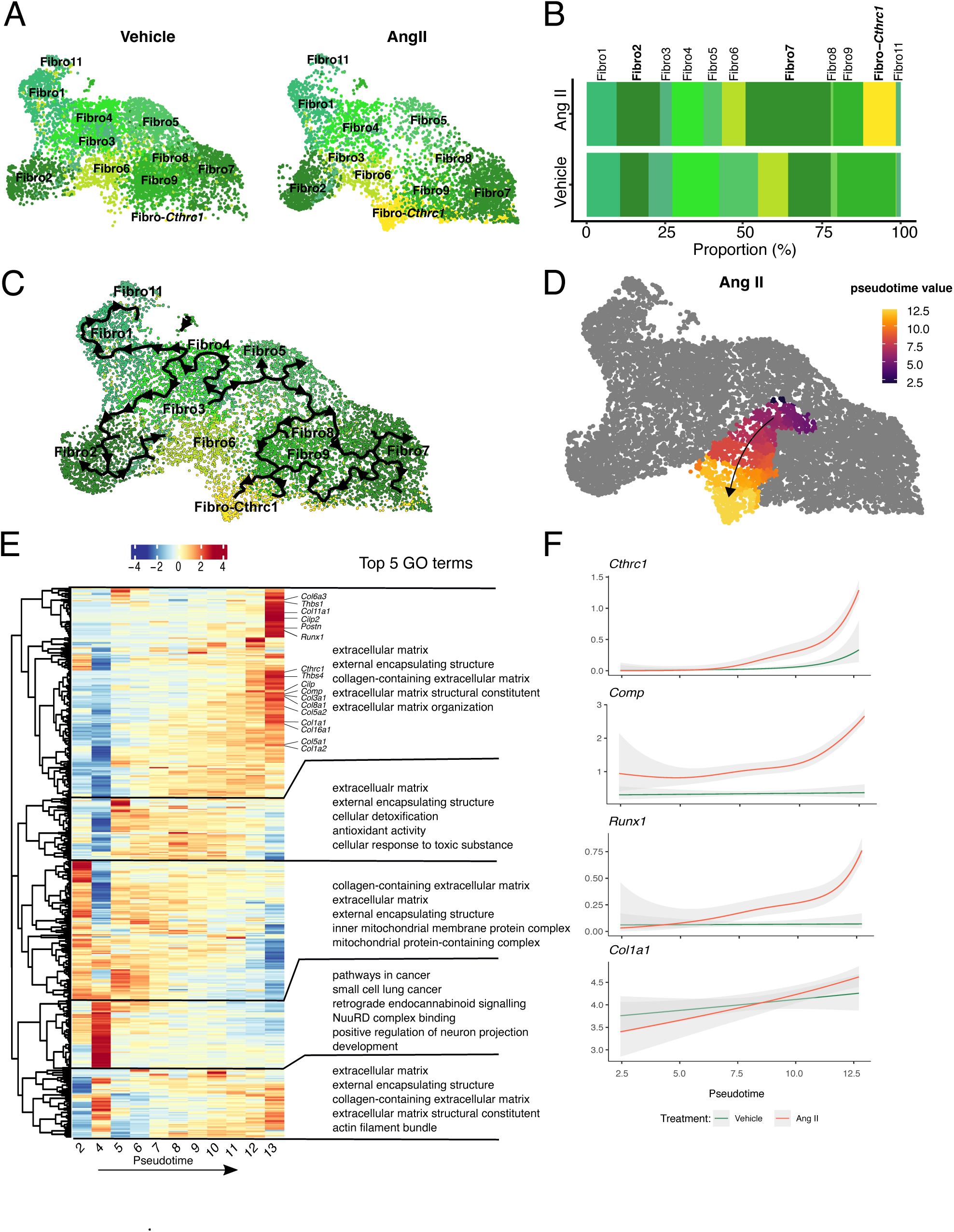
Pseudotime analysis of aortic fibroblast subclusters. UMAP projections of aortic fibroblast (fibro) subclusters from vehicle- and angiotensin II (Ang II)-treated mice (A). Stacked percentage bar graphs summarizing the relative proportions of cells belong to different fibroblast sub populations in normotensive and hypertensive mouse aortas. Width of the individual boxes comprising each bar represent the proportion of cells classified as each fibroblast cell population (B). Trajectory graph developed for fibroblast subclusters from Monocle3 analysis. The black lines show the structure of the graph (C). Supervised trajectory selected corresponding to Fibro-*Cthrc1* development estimated from Monocle3. Colour indicates the calculated pseudotime values and black arrow indicate the direction of the trajectory (D). Heatmap of top 500 genes with a changing expression pattern over calculated pseudotime in D in Ang II-treated cells and top 5 GO pathways enriched for each gene cluster (E). The x-axis represents the pseudotime, while the colour (red for high expression, blue for low expression) indicates the varying levels of gene expression. The heatmap was clustered into five gene clusters by hierarchical clustering considering changing expression patterns over pseudotime. Smoothed gene expression patterns for *Cthrc1*, *Comp*, *Runx1*, *Col1a1* and *Scx* over pseudotime (x-axis) separated by condition (green-vehicle, red-Ang II, grey error band-95% confidence interval).

To investigate potential differentiation pathways between fibroblast subclusters, we performed RNA trajectory analysis (Figure 6C). Pseudotime trajectory analysis across all fibroblast subclusters suggests Fibro-*Cthrc1* (and other fibroblast subsets) develop from Fibro9 cells (Figure 6D; Supplementary Table 3). To better understand the gene expression changes involved in the potential differentiation of Fibro9 to Fibro-*Cthrc1* we focused on this specific branch of the differentiation trajectory (Figure 6D). To identify genes that significantly changed over this trajectory, we sorted genes based on their expression profiles along the pseudotime (Figure 6E, Supplementary Table 13). Examining genes enriched across the differentiation trajectory time points in angiotensin II-infused mice only, we observed a distinct group of genes with higher expression towards the end of the trajectory which were predicted to be associated with ECM remodelling (i.e., *extracellular matrix*; *external encapsulating structure*; *collagen-containing extracellular matrix*; *extracellular matrix structural constituent and extracellular matrix organization*; Figure 6E). Additionally, upon analysing individual genes, we found that the expression of *Cthrc1*, *Comp*, and *Runx1* increased along the differentiation trajectory, with significantly higher expression in angiotensin II-infused mice compared to vehicle-infused (Figure 6F). Conversely, *Col1a1* increased along the differentiation trajectory, but to a similar extent in vehicle and angiotensin II-infused mice. These findings suggest a substantial change in the ECM-related expression profile of Fibro9 during its potential differentiation into Fibro-*Cthrc1*.

Next, we sought to determine distinct features of Fibro-*Cthrc1*. GO enrichment analysis revealed that Fibro-*Cthrc1* is highly profibrotic compared to all other fibroblast subclusters in vehicle- and angiotensin II-infused mice with numerous upregulated ECM remodelling GO terms (Figure 7A-B). Furthermore, transcripts for collagens (*Col1a1*, *Col1a2*, *Col3a1*, *Col5a2, Col8a1* and *Col11a1*), elastin (*Eln*) and the matricellular protein periostin (*Postn*) were significantly upregulated in Fibro-*Cthrc1* compared to all other aortic fibroblasts in angiotensin II-infused mice, suggesting they are the most powerful fibrogenic cell population in our dataset (Figure 7C). This was further supported by Fibro-*Cthrc1* displaying the highest expression of genes related to *ECM organisation*, *extracellular fibril organisation* and *collagen fibril organisation* GO terms compared to all other fibroblasts (Figure 7D). Interestingly, *Acta2* – the classical marker for myofibroblasts – was not highly expressed in Fibro-*Cthrc1*. Rather, *Acta2* expression was most prominent in Fibro7, suggesting this subcluster may represent classical myofibroblasts (Figure 7E).

**Figure 7.**
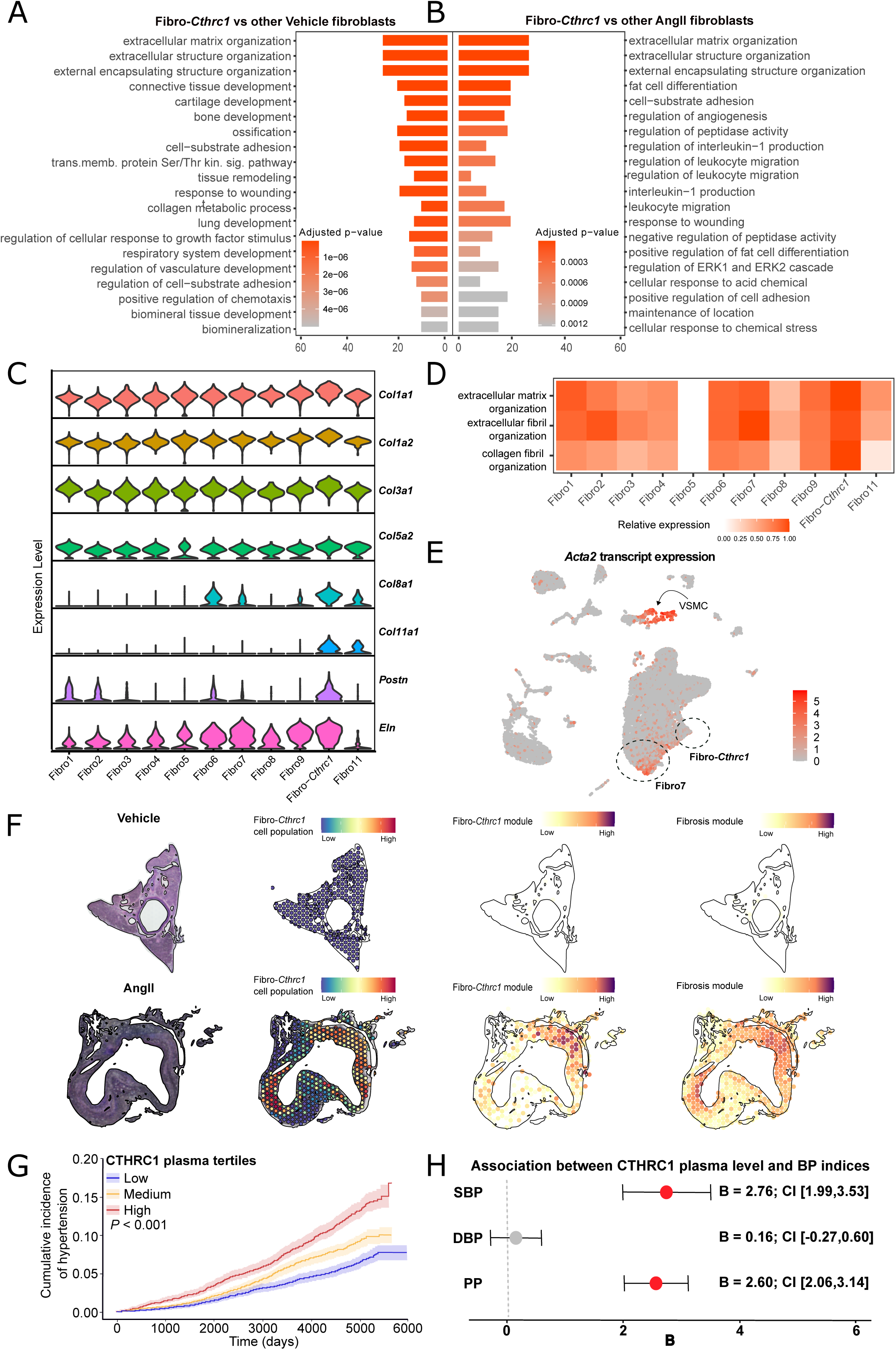
Profibrotic genes are highly enriched in Fibro-*Cthrc1* in aortic stiffening. GO terms enriched in a set of genes derived from comparing gene expression differences between Fibro-*Cthrc1* vs all other fibroblast types in aortas from vehicle-(A) and angiotensin II (Ang II)-treated mice (B) the number of genes mapped to each GO term, and colour indicates the adjusted *P* value from GO enrichment analysis. Violin plot projections of *Col1a1*, *Col1a2*, *Col3a1, Col5a2*, *Col8a1*, *Col11a1, Postn* and *Eln* gene expression among fibroblast subclusters from Ang II-treated mice. Shape of the coloured sections represent the probability distribution of the gene expression levels where wider sections represent higher probability, and narrower sections represent lower probability. Heat map showing relative total transcript expression of fibroblast genes classified within the GO categories shown on y axis. Colour represents the sum of transcripts for genes corresponding to each GO term within each cell population (red = high, white = no/low; D). UMAP plot with cells coloured according to *Acta2* transcript abundance (red = high, grey = low; E). Localisation of the Fibro-*Cthrc1* cell population in aortas from vehicle-(F, top panel) and Ang II-treated mice (F, bottom panel) using spatial transcriptomics integrated with scRNA-seq. Colour scale is relative to average expression of module genes for module genes. Cumulative incidence of hypertension in different plasma tertiles of CTHRC1 (analysis performed in 13,272 normotensive subjects; no BP-lowering medications, BP<140/90 mmHg with no previous diagnosis of hypertension; G). Beta values for general linear models adjusted for age, sex, BMI, smoking status and alcohol intake frequency in 23,564 normotensive subjects (no BP-lowering medications, BP<140/90 mmHg with no previous diagnosis of hypertension; H). Fibro, fibroblast; Thr, threonine; transmemb, transmembrane; Ser, serine; sig, signalling; stim, stimulation; VSMC, vascular smooth muscle cells.

Following integration with the aortic cellulome dataset, spatial transcriptomics confirmed that Fibro-*Cthrc1* cells were only present in aortas of angiotensin II-infused, and not vehicle-infused mice (Figure 7F). Spatial transcriptomics confirmed that the specific regions of the vessels exhibiting gene signatures corresponding to Fibro-*Cthrc1* (“Fibro-*Cthrc1*” module: *Comp*, *Cthrc1*, *Col11a1*, *Kcnma1*, *Ddha1*, *Thbs2* and *Col8a2;* Supplementary Figure 7), also expressed the gene signatures of fibrosis (“fibrosis module”: *Postn*, *Sparc*, *Dcn*, *Fn1*, *Fbln5*, *Ccn2*, *Col1a1*, *Col2a1*, *Col3a1*, *Col5a2*; Figure 7F, Supplementary Figure 8). Neither module was present in aortas from normotensive mice (Figure 7F). While the Fibro-*Cthrc1* and fibrosis modules were not observed in aortas from normotensive mice, some of the individual genes of each module were present, including *Comp* from the Fibro-*Cthrc1* module (Supplementary Figure 7) and *Postn*, *Sparc*, *Dcn*, *Fbln5*, *Col1a1*, *Col3a1*, and *Col5a2* from the fibrosis module (Supplementary Figure 8). All individual genes from the Fibro-*Cthrc1* and fibrosis modules were enriched in aortas from angiotensin II-infused mice (Supplementary Figures 7 & 8).

### 3.7 High plasma CTHRC1 levels are associated with increased risk of hypertension

Considering that *Cthrc1* expression is upregulated in the aortic cellulome, and in other cardiovascular tissues after Ang II treatment^37, 38^, we explored the potential for CTHRC1 to be used as a marker/predictor for the development of hypertension. A normotensive sub-population within UK Biobank Pharma Proteomics Project was divided into tertiles based on the baseline plasma levels of CTHRC1. The tertile with highest plasma CTHRC1 levels showed the highest cumulative incidence of hypertension over approximately 15 years of follow up (Figure 7G). This tertile also had the most unfavourable cardiovascular profile (except for alcohol intake frequency, Table 1). The differences in baseline characteristics may explain why the highest tertile shows an increased risk of hypertension. Indeed, when the Cox regression is not adjusted for confounders, CTHRC1 is associated with an increased hazard ratio of hypertension (HR = 3.01; 95% CI [2.55-3.56]; *P* < 0.001). This association is abolished when the Cox regression is adjusted for some hypertension risk factors like age, sex, BMI, smoking and alcohol (HR = 1.20; 95% CI [0.98-1.46]; *P* = 0.08).

In a general linear model adjusted for age, sex, BMI, smoking status and alcohol intake frequency, CTHRC1 plasma levels were significantly associated with SBP (B = 2.76; 95% CI [1.99-3.53]; *P* < 0.001) and PP (B = 2.60; 95% CI [2.06-3.14]; *P* < 0.001), but not DBP (B = 0.16; 95% CI [-0.27-0.60]; *P* = 0.46; Figure 7H). Collectively, these data indicate that baseline plasma CTHRC1 levels are associated with baseline SBP and PP. Moreover, individuals with high plasma levels of CTHRC1 have an unfavourable cardiovascular profile and an increased risk of hypertension.

## 4. Discussion

Here, we have comprehensively mapped the aortic cellulome in the setting of hypertension, to provide insights into the cellular and molecular drivers of aortic pathological remodelling and fibrosis. All major cell types in the aorta had detectable transcriptional responses to hypertension, with *extracellular matrix organization* gene pathways highly enriched in fibroblasts. A key difference between hypertensive versus normotensive aortas was the presence of a distinct fibroblast subcluster (termed Fibro-*Cthrc1*) in the former. Fibro-*Cthrc1* were the most fibrogenic cell type detected in our dataset, suggesting they may be key drivers of aortic stiffening in hypertension.

Previous studies have used single-cell transcriptomics to map the aortic cellulome in various disease settings such as atherosclerosis^39–41^, obesity^6^, Marfan syndrome^8^, and abdominal aortic aneurysms^7^. While hypertension may play a role in some of these disease settings, no studies to date have reported on the impact of hypertension alone on the aortic cellulome. Some studies focused on specific cell types such as endothelial or immune cells^42, 43^, whereas others mapped the whole aortic cellulome^6–8^. Indeed, we report similar cellular subclusters as other studies that analysed the whole aortic cellulome, particularly with respect to classical vascular cells such as endothelial and VSMCs^6–8^. A distinguishing feature of our protocol was the isolation of live, metabolically active nucleated cells prior to scRNA-seq library construction^14, 15^, ensuring that a higher proportion of high-quality cells were sequenced (reducing the number of cells that are later excluded during data processing). In the present study, fibroblasts were by far the most abundant and heterogeneous cell type within the aortic dataset. This contrasts other scRNA-seq studies on whole aortas where immune cells were more abundant than fibroblasts^6, 8^. This discrepancy between studies is possibly due to differences in the digestion protocol used to isolate single cells or the disease states being studied. Moreover, it highlights the limitation in interpreting cell-type abundances in scRNA-seq datasets, and the importance of using other techniques (such as spatial transcriptomics, histopathology and IHC) to validate interpretations from scRNA-seq datasets.

While hypertension was associated with marked changes in gene expression in all cell types, fibroblasts were the leading cell type to adopt a transcription profile aligned with enhanced ECM remodelling, as determined by GO analysis. Notably, endothelial cells and VSMCs were the only other cell types with upregulated ECM remodelling GO terms in response to Ang II. Despite previous reports that macrophages are major contributors to vascular stiffening in hypertension, ECM remodelling pathways were not upregulated in macrophages or indeed any of the other major immune cell populations identified by scRNA-seq in the present study. Rather, hypertension was associated with upregulation of inflammation pathways in immune cell subsets (i.e. immune cell activation, differentiation and proliferation). These findings are consistent with a previous aortic scRNA-seq study which identified proinflammatory aortic macrophage phenotypes in atherosclerosis^39^. Other major immune cell subclusters (such as dendritic cells and lymphocytes) have not been well characterised in previous aortic scRNA-seq studies. While not the focus of the current study, future scRNA-seq studies interrogating the role of aortic leukocytes in hypertension within the current and other available datasets would be insightful.

Among the fibroblast subtypes detected, one subpopulation – Fibro-*Cthrc1* – stood out due to its prominence in hypertensive aortas and complete absence in healthy blood vessels. A distinctive feature of this fibroblast subtype was it high levels of expression of *Cthrc1* transcripts, which encode for the secreted glycoprotein Collagen Triple Helix Repeat Containing 1 (CTHRC1). CTHRC1 was first discovered 20 years ago in a screen for differentially expressed proteins in balloon-injured versus normal rat arteries^44^. CTHRC1 is a secreted 28-kDa protein that is glycosylated and highly conserved across species^38^. Although CTHRC1 is typically found in areas of high interstitial collagen deposition and TGF-β activity, its role in fibrosis is somewhat contentious^45^. On the one hand, overexpression of CTHRC1 in fibroblasts and vascular smooth muscle cells (VSMCs) inhibits collagen synthesis and promotes collagen degradation. It also terminates TGF-β signalling by reducing SMAD2/3 phosphorylation. However, despite these apparent anti-fibrotic actions, numerous studies in various pathological conditions have attributed a pro-fibrotic function to CTHRC1-expressing fibroblasts^45^. In the present study, Fibro-*Cthrc1* expressed high levels of other transcripts with known roles in matrix remodelling and fibrosis, including collagen genes and periostin. GO analysis identified Fibro-*Cthrc1* as the most pro-fibrotic of all cell types, including other fibroblasts, in the hypertensive aorta. Previously, we identified an analogous population of fibroblasts in the hearts of angiotensin II-treated mice^37^. These thrombospondin 4-expressing fibroblasts (Fibroblast-*Thbs4*) expressed high levels of *Cthrc1*, along with other genes involved in ECM remodelling, such as *Fmod* and *Cilp.* Fibroblast-*Thbs4* cells were aggregated within fibrotic foci but notably absent from perivascular fibrotic lesions, suggesting their specific involvement in interstitial fibrosis^37^. Similarly, a fibroblast subcluster with high *Cthrc1* expression was identified in diseased human lungs and mouse models of lung fibrosis. These fibroblasts were only present in diseased lung tissue and displayed high expression of several collagen genes, indicative of a highly fibrogenic phenotype^46^. Finally, scRNA-seq analysis of hearts post-myocardial infarction identified a *Cthrc1*-expressing fibroblast subcluster with a pro-fibrotic signature that localized to the scar. Hence, the identification of Fibro-*Cthrc1* in the present study adds to the growing body of evidence that these cells are crucial drivers of fibrosis under disease conditions^47^.

There is a prevailing paradigm that myofibroblasts – which differentiate from fibroblasts in the presence of cytokines and trophic factors such as transforming growth factor β - are the main protagonists in driving pathological fibrosis^4, 5^. Indeed, we observed an increase in a population of fibroblasts expressing the myofibroblast marker, *Acta2*, in aortas from angiotensin II-infused mice (defined as Fibro7 subcluster). Most strikingly, these cells were clearly a distinct population from Fibro-*Cthrc1* which our trajectory inference analysis suggests also emerges from Fibro9. Furthermore, GO enrichment analysis indicated that *Acta2*-expressing Fibro7 have far lower fibrogenic capacity that Fibro-*Cthrc1*. Hence, our study challenges the current paradigm by suggesting that Fibro-*Cthrc1*, rather than myofibroblasts, are the main cellular drivers of aortic fibrosis in hypertension^46,47^.

Our spatial transcriptomics data confirm that the Fibro-*Cthrc1* subcluster was only detected in aortas from angiotensin II-infused mice. This was further validated with immunohistochemistry showing CTHRC1 was localised to the adventitia of angiotensin II-but not vehicle-infused mouse aortas. Immunohistochemistry also showed that CTHRC1 was localised to the PVAT of the aorta; however, this did not differ between vehicle- and angiotensin II-infused mice. The absence of Fibro-*Cthrc1* module detection in vehicle-infused mice suggests that CTHRC1 was expressed by cell types not included in the scRNA-seq dataset. Indeed, previous studies have shown that adipocytes can secrete CTHRC1^48^. Although the PVAT was included in the scRNA-seq cell suspension, adipocytes were not represented in the final dataset as they were excluded at the cell sorting stage of sample preparation due their large cell size.

Finally, we investigated the link between circulating CTHRC1 and hypertension using the UK Biobank. High plasma CTHRC1 levels in normotensive patients were linked to an unfavourable cardiovascular profile and a higher risk of developing hypertension over 15 years. CTHRC1 levels were also associated with higher SBP and PP, which are influenced by large artery stiffening. Previous studies have shown that increased circulating CTHRC1 levels are linked to other chronic conditions, such as rheumatoid arthritis, where CTHRC1 acts as a pathogenic driver. In rheumatoid arthritis, plasma CTHRC1 levels are higher compared to osteoarthritis patients and healthy individuals, correlating with disease severity^49^. Similarly, a smaller-scale investigation in 44 patients with or without chronic kidney disease – a condition also known to be associated with tissue fibrosis – showed that plasma CTHRC1 levels were correlated with increased proteinuria, plasma creatinine and cholesterol^50^. Hence, collectively these studies suggest that circulating CTHRC1 may be a useful biomarker for chronic diseases associated with fibrosis and inflammation.

In conclusion, the present study identifies a unique fibroblast population (Fibro-*Cthrc1*) in the hypertensive mouse aorta with phenotypic properties that suggest it is likely a pathogenic mediator of aortic fibrosis and stiffening. Our study challenges the current paradigm by suggesting that Fibro-*Cthrc1*, as opposed to myofibroblasts, are the main drivers of fibrosis and vascular stiffening in hypertension. Our findings provide a foundation for further characterisation of the pathobiology of Fibro-*Cthrc1* in hypertension and their validation as a potential therapeutic and/or diagnostic target.

## 6. Author contribution statement

M.J., M.I.D, A.R.P., A.V. and G.R.D contributed to the conception and design of the work; M.J., T.G.H., G.F., A.H., V.T., H.D., Q.N.D., T.G., M.G.L., A.R.P, T.J.G., M.L. and A.V. contributed to the acquisition of the data. M.J., M.I.D., R.H., T.G.H., R.C., V.T., I.H., T.J.G., M.L. A.R.P., A.V. and G.R.D. contributed to the analysis and interpretation of the data. M.J., M.I.D., A.R.P, A.V. and G.R.D contributed to the drafting the manuscript; and all authors approved the submission of this work.

## 7. Acknowledgments

The authors thank Dr Bhupinder Pal and David Baloyan for their assistance with the tissue preparation and 10X Genomics experiments and the La Trobe Animal Research and Teaching Facility staff for their assistance with the animal husbandry.

