## Supplementary data for "High-resolution transcriptomic profiling of the aortic cellular landscape during hypertension reveals novel drivers of vascular fibrosis"

**Short title:** Novel cellular drivers of aortic stiffening in hypertension

**Corresponding authors:**	A/Prof Alexander R Pinto, PhD

Baker Heart and Diabetes Institute

75 Commercial Road

Prahran, Victoria, Australia, 3004

Ph: +61 3 8532 1275

A/Prof Antony Vinh

Prof Grant Drummond

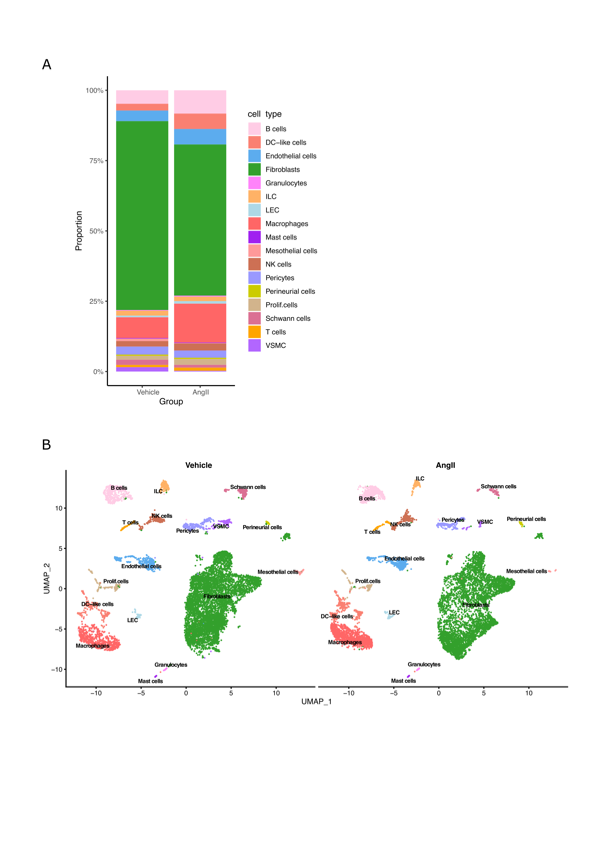

**Supplementary Figure 1. Relative proportions of aortic cell populations captured from scRNA-seq across two experimental groups.**

Stacked percentage bar graph summarizing the relative proportions of cell types analysed from individual samples from normotensive and hypertensive mouse aorta. Heights of the individual boxes comprising each bar represent the proportion of cells classified as each cell population (A). UMAP projection of aortic cells analysed by scRNAseq in both vehicle- and Ang II-treated mice (B). Each dot represents a cell and cells are coloured by distinct cell populations as indicate (n = 3, per group).
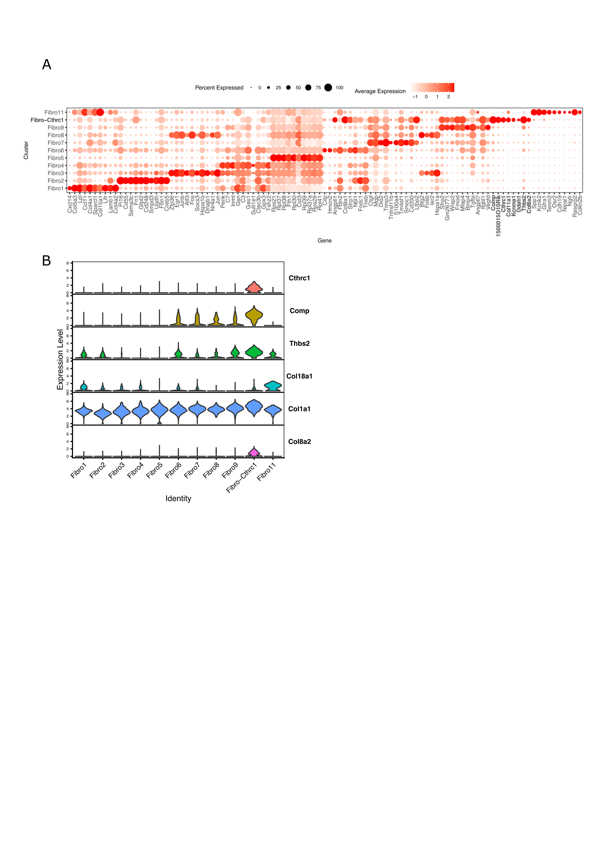

**Supplementary Figure 2. Marker genes for newly emerged fibroblast cell population.**

Dot plot summarizing the expression of top10 marker genes identified for each fibroblasts sub-population (A). Dot colour and size indicate the relative average expression level and the proportion of cells expressing the gene, respectively, within each cell population. Violin plot projections of *Cthrc1*, *Comp*, *Thbs2,* *Col18a1*, *Col1a1* and *Col8a2* gene expression among fibroblast sup-populations.

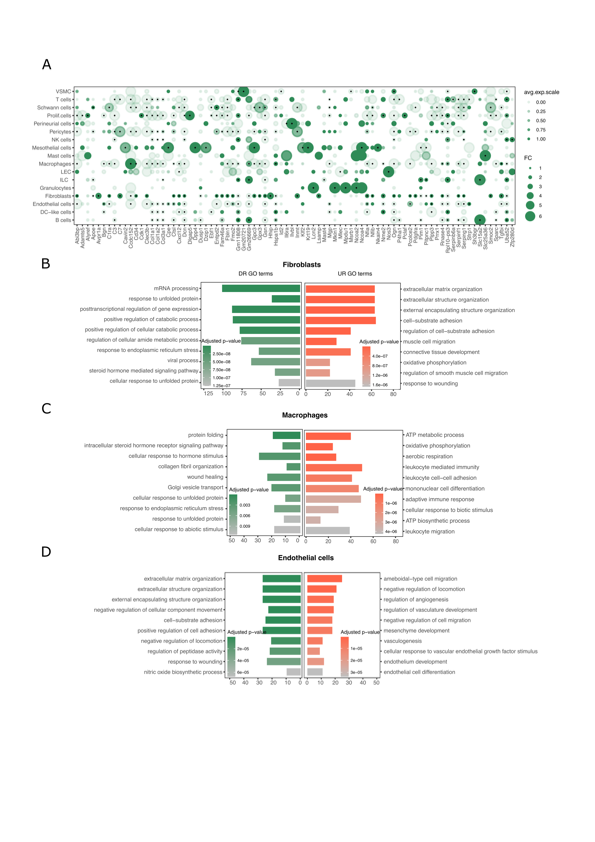

**Supplementary Figure 3. Gene expression changes in aortic cell populations.**

Dot plot summarizing the expression of genes identified as within the top 10 genes downregulated (uncorrected *P*<0.001 and FC>2) in response to Ang II administration, for each cell type (A). Dot colour and size are proportional to average expression within each cell cluster in control cells and fold change compared to the Ang II cells, respectively. The black dots in the centre of some circles indicate the statistical significance of *P*<0.01. Combined GO bar plots summarizing top 10 up (red) and down (green)-regulated GO pathways enriched by up and down regulated genes in fibroblasts (B) macrophages (C) and endothelial cells (D) in response to Ang II treatment. *x* axis indicates the number of genes mapped to each GO term, and colour indicates the adjusted *P* value from GO enrichment analysis. GO terms are ordered based on statistical significance.

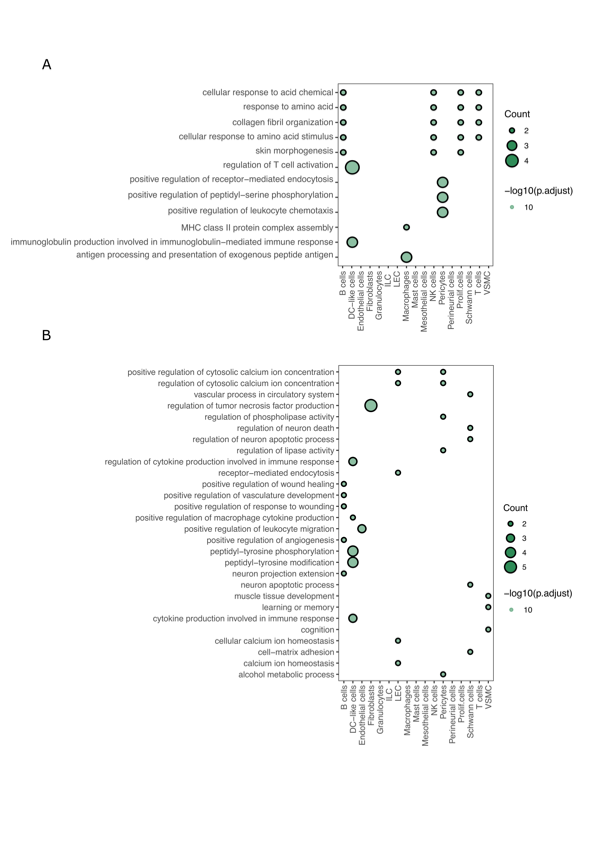

**Supplementary Figure 4. Gene ontology pathways enriched by downregulated ligand/ receptor genes**

Dot plot summarizing top 5 GO terms enriched in a set of genes that encode ligands (A) and receptors (B) downregulated after Ang II treatment. GO terms are ordered by their frequency of significant enrichment in different aortic cell populations. Dot colour and size proportional to the statistical significance and number of down regulated genes enriched in each GO term, respectively.

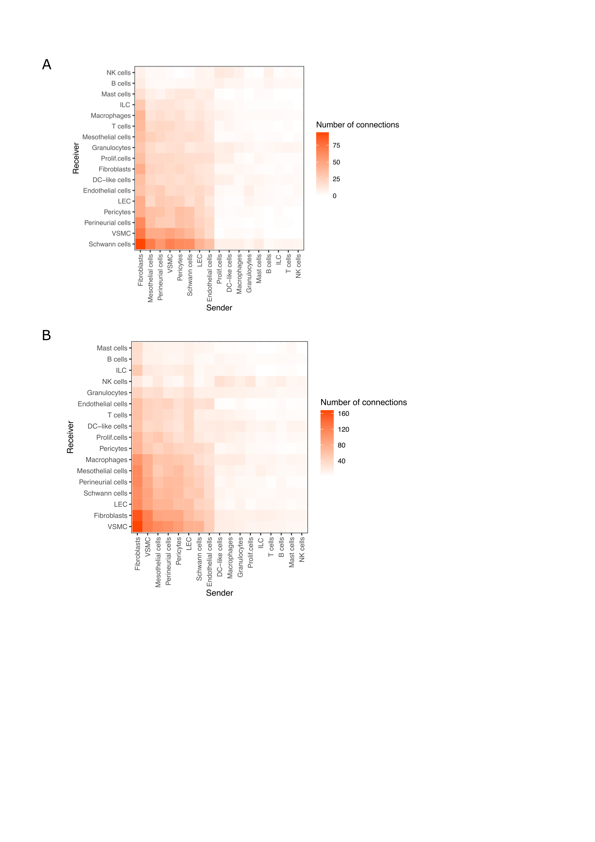

**Supplementary Figure 5. Interactions between aortic cell populations in vehicle and Ang II-treated mice**

Heatmaps summarizing number of intercellular connections between aortic cell populations in vehicle- (A) and Ang II-treated (B) mice. The x-axis represents the sender cell type, which releases the signaling ligand, while the y-axis represents the recipient cell type expressing the corresponding receptor(s) to receive signals. Each square is color-coded based on the number of connections, with red indicating high connectivity and white indicating low connectivity. Cell populations are ordered based on number of interactions to identify most interactive aortic cell populations within the aortic intercellular network.

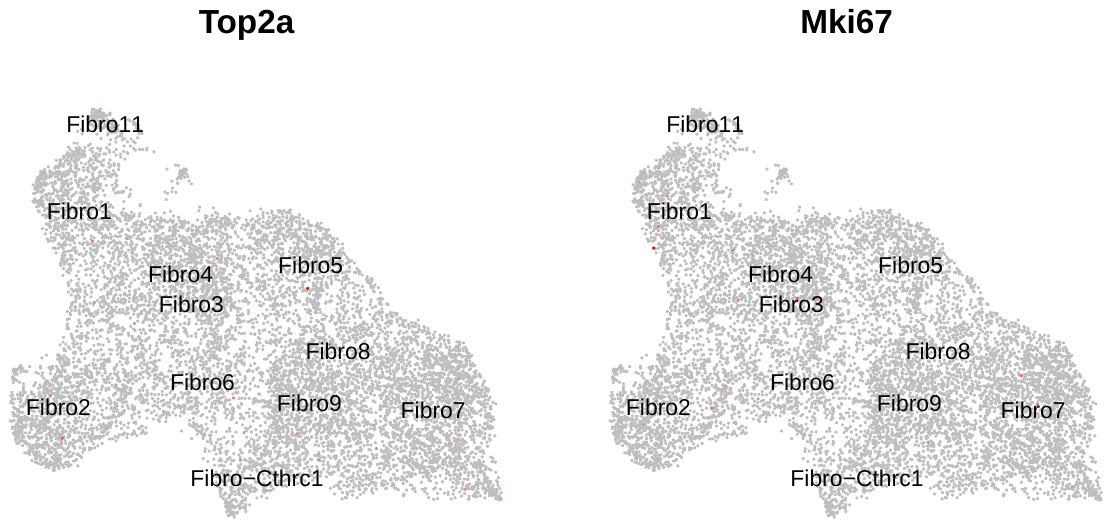

**Supplementary Figure 6. Expression of genes marking cell proliferation**

Feature UMAP projections of cell proliferation marker genes Top2a and Mki67 on aortic fibroblast cell populations. Each dot represents a cell and cells are coloured based on the relative gene expression (red=high; grey=low/no gene expression).

**Supplementary Figure 7: Spatial gene expression of Fibro-*Cthrc1* module genes in vehicle- and Ang II-treated mouse aorta.**

Representative images of spatial feature plot of *Cthrc1* module genes *Comp*, *Cthrc1*, *Col11a1*, *Kcnma1*, *Ddah1 Thbs2* and  *Col8a2* in aorta from vehicle- (A) and Ang II-treated (B) mice. Colour scale is relative to average expression.

**Supplementary Figure 8: Spatial gene expression of fibrosis module genes in vehicle- and Ang II-treated mouse aorta.**

Representative images of spatial feature plot of fibrosis module genes *Postn, Sparc, Dcn, Fbln5, Fn1, cn2, Col1a1, Col2a1, Col3a1* and *Col5a2* in aorta from vehicle- (A) and Ang II-treated (B) mice. Colour scale is relative to average expression.
